# Early biomarker discovery in urine of Walker 256 subcutaneous rat model

**DOI:** 10.1101/114611

**Authors:** Jianqiang Wu, Zhengguang Guo, Youhe Gao

## Abstract

Despite advances in cancer treatments, early detection of cancer is still the most promising way to improve outcomes. Without homeostatic control, urine reflects early changes in the body and can potentially be used for early cancer diagnosis. In this study, a Walker 256 tumor rat model was established by subcutaneous injection of Walker 256 tumor cells. To identify urinary proteome changes during cancer development, urine samples from Walker 256 tumor-bearing rats were collected at five time points corresponding to before cancer cell implantation, before tumor mass palpability, at tumor mass appearance, during rapid tumor growth, and at cachexia. The urinary protein patterns changed significantly as the tumors progressed, as measured using sodium dodecyl sulfate-polyacrylamide gel electrophoresis (SDS-PAGE). The urinary proteome of tumor-bearing rats was identified using Fusion Lumos mass spectrometry with label-free quantification. Then, 30 dynamically changed urinary proteins during cancer progression were selected as more reliable cancer biomarkers, and they were validated by targeted proteomics. Combined with the results of label-free and targeted proteome quantification, a total of 10 urinary proteins (HPT, APOA4, CO4, B2MG, A1AG, CATC, VCAM1, CALB1, CSPG4, and VTDB) changed significantly even before a tumor mass was palpable, and these early changes in urine could also be identified with differential abundance at late stages of cancer. Our study indicated that urine is a sensitive biomarker source for early detection of cancer.

## Introduction

Cancer is an important public health concern worldwide and is the second leading cause of death in the United States [1]. The early detection of in situ or invasive carcinoma may prevent cancerous metastatic processes; thus, early detection can significantly improve survival rates for cancer patients. Despite technical advances in tumor diagnosis in the last decade, there are still many cancer patients who cannot be diagnosed at early disease stages. To reduce mortality from cancer, novel approaches must be considered for screening and early detection of cancer.

Cancer biomarkers are measurable changes associated with the pathophysiological processes of cancers that have the potential to diagnosis cancer and monitor cancer progression. Urine is a promising bio-fluid for biomarker research. Unlike blood, urine has no mechanisms to maintain a homeostatic state and can reflect systemic changes in the body. As a sensitive biomarker sample source, urine has the potential to reflect pathological changes, especially in the early phase of disease [2, 3]. In recent years, advances in proteomics, especially in mass spectrometry, have led to the identification of more than 3000 unique proteins in human urine. Meanwhile, urinary proteomics has been successfully applied to discover novel biomarkers for cancer diagnosis and cancer monitoring [4-6]. However, whether urine protein biomarkers assist in the early diagnosis of cancer is unclear.

The Walker 256 (W256) tumor-bearing rat model is a classic animal model with which to study tumor progression and tumor cachexia. In this study, a tumor-bearing rat model was established by subcutaneous injection of W256 tumor cells. To identify changes in the urinary proteome during cancer development, urine samples from tumor-bearing rats were collected at five time points corresponding to before cancer cell implantation, before the tumor mass was palpable, tumor mass appearance, rapid tumor growth, and cachexia. Using label-free proteomics analysis and multiple reaction monitoring (MRM)-based validation, cancer-associated urine biomarkers were identified.

## Materials and methods

### Animals

Male Wistar rats (150 ± 20 g) were supplied by the Institute of Laboratory Animal Science, Chinese Academy of Medical Science. All animals were maintained with free access to a standard laboratory diet and water with a 12-h light-dark cycle under controlled indoor temperature (22 ± 2°C) and humidity (65–70%) conditions. Animal procedures were approved by the Institute of Basic Medical Sciences Animal Ethics Committee, Peking Union Medical College (ID: ACUC-A02-2014-007), and the study was performed according to guidelines developed by the Institutional Animal Care and Use Committee of Peking Union Medical College.

### Tumor model

A subcutaneous tumor-bearing animal model was established as previously reported [7]. Walker-256 (W256) carcinosarcoma cells were implanted intraperitoneally into Wistar rats. Seven days following implantation, the ascitic tumor cells were harvested from the peritoneal cavity. W256 tumor cells used for establishing the animal model were obtained from the ascitic fluid after two cell passages. Then, W256 cells were collected, centrifuged, and resuspended in phosphate-buffered saline (PBS). The viability of W256 cells was evaluated by the Trypan blue exclusion test using a Neubauer chamber.

The rats were randomly divided into two groups: tumor-bearing rats (n=10) and control rats (n=5). Tumor-bearing rats were subcutaneously inoculated with 2 × 10^6^ viable W256 cells in 0.2 mL of PBS into the right flank of the animal. An equal volume of PBS was subcutaneously inoculated into the control rats. During inoculation procedures, the animals were anesthetized with sodium pentobarbital solution (4 mg/kg).

### Urine collection and sample preparation

After the rats were acclimated in metabolic cages for 3 days, urine samples were collected from each rat on days 0, 4, 6, 9, 11, and 14 after cell or PBS inoculation. Animals were individually placed in metabolic cages for 8 hours to collect urine samples. During urine collection, rats had free access to water but no food to avoid urine contamination.

After urine collection, urine samples were immediately centrifuged at 12,000 g for 30 min at 4°C to remove cell debris. The supernatants were precipitated with three volumes of ethanol at 4°C, followed by centrifugation at 12,000 g for 30 min. The pellet was then resuspended in lysis buffer (8 M urea, 2 M thiourea, 50 mM Tris, and 25 mM DTT). The protein concentration of each sample was measured using the Bradford assay.

### SDS-PAGE analysis

For each sample, on day 0, 4, 6, 9, 11, and 14 after W256 cell inoculation, 30 μg of protein was added to the sample loading buffer (50 mM Tris-HCl, pH 6.8, 50 mM DTT, 0.5% SDS, and 10% glycerol) and incubated at 97°C for 10 min. The proteins were then resolved by 12% sodium dodecyl sulfate-polyacrylamide gel electrophoresis (SDS-PAGE). After electrophoresis, the gels were stained using Coomassie brilliant blue. Urine samples from four randomly selected tumor-bearing rats were used for SDS-PAGE.

### Tryptic digestion

The urine samples on days 0, 4, 6, 9, and 14 of four tumor-bearing rats after W256 cell inoculation were selected for further proteomic analysis. The urinary proteins were prepared using the filter-aided sample preparation method as previously described [8]. Each 100 μg of protein was denatured with 20 mM dithiothreitol at 37°C for 1 h and alkylated with 50 mM iodoacetamide in the dark for 30 min. Then, samples were loaded onto 10-kD filter devices (Pall, Port Washington, NY, USA) and centrifuged at 14,000 g at 18°C. After washing twice with UA (8 M urea in 0.1 M Tris-HCl, pH 8.5) and four times with 25 mM NH4HCO3, the samples were digested with trypsin (enzyme to protein ratio of 1:50) at 37°C overnight. The peptide mixtures were desalted using Oasis HLB cartridges (Waters, Milford, MA) and dried by vacuum evaporation.

### Liquid chromatography coupled with tandem mass spectrometry (LC-MS/MS) analysis

The twenty peptide samples resulting from the above digestion were re-dissolved in 0.1% formic acid to a concentration of 0.5 μg/μL. For analysis, the peptides were loaded on a trap column (75 μm × 2 cm, 3 μm, C18, 100 Å) and were separated on a reverse-phase analytical column (50 μm × 150 mm, 2 μm, C18, 100 Å) using the Thermo EASY-nLC 1200 HPLC system. Then, the peptides were analyzed with a Fusion Lumos mass spectrometer (Thermo Fisher Scientific, Bremen, Germany). The elution gradient for the analytical column was 95% mobile phase A (0.1% formic acid, 99.9% water) to 40% mobile phase B (0.1% formic acid, 89.9% acetonitrile) over 120 min at a flow rate of 300 nL/min. The mass spectrometer was set in positive ion mode and operated in a data-dependent acquisition mode with a full MS scan from 150 to 2,000 m/z and MS/MS scan from 110 to 2,000 m/z with a resolution of 120,000. Dynamic exclusion was employed with a 30-s window.

### Label-free proteome quantification

The raw files of proteomic data were searched against the SwissProt rat database (released in July 2016, containing 7,973 sequences) using Mascot software (version 2.5.1, Matrix Science, London, UK). The parent ion tolerance was set to 10 ppm, and the fragment ion mass tolerance was set to 0.05 Da. Carbamidomethyl of cysteine was set as a fixed modification, and the oxidation of methionine was considered a variable modification. The specificity of trypsin digestion was set for cleavage after K or R, and two missed trypsin cleavage sites were allowed. Peptide and protein identification was further validated using Scaffold (version 4.4.0, Proteome Software Inc., Portland, OR). Peptide identifications were accepted at an FDR less than 1.0% by the Scaffold Local FDR algorithm, and protein identifications were accepted at an FDR less than 1.0% with at least 2 unique peptides. Comparisons across different samples were performed after normalization of total spectra accounts using Scaffold software.

## MRM

MRM was performed on a QTRAP-6500 mass spectrometer (AB SCIEX, Framingham, MA, USA) equipped with a nano-UPLC system (Waters, Milford, MA). The peptides were eluted with 5-30% buffer B (0.1% formic acid, 99.9% ACN) at 300 nL/min for 60 min.

The raw files of MS data acquired at the biomarker screening phase were used as the MS/MS spectral library to select peptides and transitions for the MRM assays. Mascot results and the list of targeted proteins were imported into Skyline software (version 3.6) to select the most intense peptide transitions. Then, a total of 120 μg of peptides mixed from each validated sample was analyzed by mass spectrometry (MS) to further select peptides and transitions for MRM validation of targeted proteins. Individual urine samples from another four tumor-bearing rats on days 0, 4, 6, 9, and 14 were analyzed by MRM assays. Each sample has three technical duplications. Unique peptides for each protein and 4-5 transitions per peptide were used for quantification. The length of a peptide candidate was 6–25 amino acids. MRM results were analyzed using the instructions from the Skyline software [9].

## Results

### Body weight and tumor mass in Walker 256 tumor-bearing rats

From 6 days after W256 cell subcutaneous implantation, the average body weight of the tumor-bearing rats was lower than that of the control rats (**Fig 1**), and reduced food intake was observed in tumor-bearing rats. On day 9 after W256 cell inoculation, the body weight of tumor-bearing rats was significantly reduced compared with their body weights at other time points.

**Fig 1.**
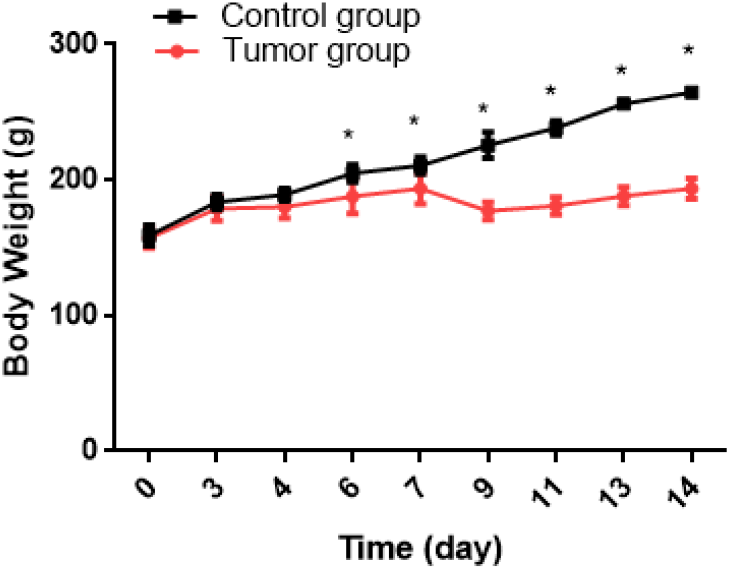
Body weights of Walker 256 tumor-bearing rats. The average body weight of the tumor group was significantly lower than that of the control group (n=10 rats in the tumor group and n=5 rats in the control group; * indicates p < 0.01). There were five time points on days 0, 4, 6, 9, and 14 corresponding to before cancer cell implantation, before the tumor mass was palpable, tumor mass appearance, rapid tumor growth, and cachexia, respectively.

The growth of a subcutaneous tumor mass in tumor-bearing rats was observed every day after W256 cell inoculation. Small tumor masses could be felt in the W256 rats beginning on the sixth day, and the tumor masses grew gradually. When the rats were sacrificed after 15 days, the tumor tissues were harvested and stained with hematoxylin and eosin (H&E) for pathological examination. Large numbers of tumor cells were observed in the tumor masses (**S1 Fig**).

### SDS-PAGE analysis of the urine samples in tumor-bearing rats

Urine samples from tumor-bearing rats collected on different days were separated by 12% SDS-PAGE. As shown in **Fig 2,** the protein patterns of urine samples in a representative tumor-bearing rat changed significantly as the tumor progressed (day 0, day 4, day 6, day 9, day 11, and day 14). Compared with the protein band on day 0, the protein bands were most significantly different on day 9 and slightly recovered on day 14. Similar patterns were observed in other rats, suggesting relatively good consistency in tumor progression. To identify changes in the urinary proteome across the development of cancer, urine samples at five time points, corresponding to before tumor cell implantation (day 0), before the tumor mass was palpable (day 4), tumor mass appearance (day 6), rapid tumor growth (day 9), and cachexia (day 14), were selected for proteomics analysis.

**Fig 2.**
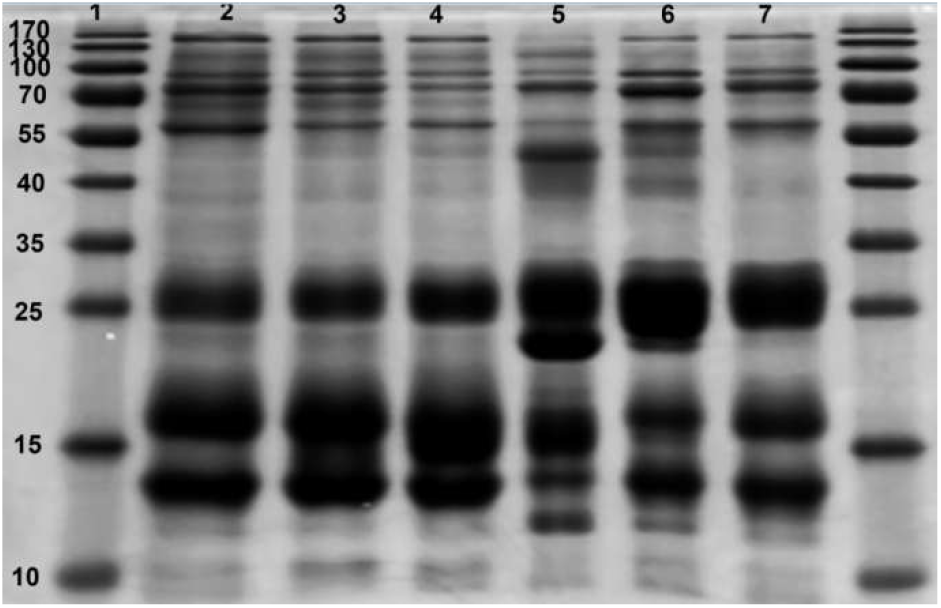
Dynamic changes in protein patterns in the urine of tumor-bearing rats. This figure is a representative diagram of tumor-bearing rats. Lane 1: Marker. Lanes 2–7, urinary proteins on day 0 (lane 2), day 4 (lane 3), day 6 (lane 4), day 9 (lane 5), day 11 (lane 6), and day 14 (lane 7) after tumor cell inoculation, respectively.

### Identification of the urine proteome with tumor progression

The study design for proteomic analysis in this study is shown in **Fig 3**. At the biomarker discovery phase, label-free LC-MS/MS quantification was used to characterize the differential expression of urinary proteins at various tumor progression stages. To investigate changes in the urine proteome with tumor progression, urine samples at 5 time points from 4 tumor-bearing rats were analyzed. A total of 20 samples were analyzed using a Fusion Lumos mass spectrometry platform. All spectra were searched against the SwissProt database using Mascot software, and the resulting data were validated using Scaffold software for protein identification and quantification. A total of 533 urinary proteins with at least 2 unique peptides were identified with 1% FDR at the protein level. All identification and quantification details are presented in **S1 Table**.

**Fig 3.**
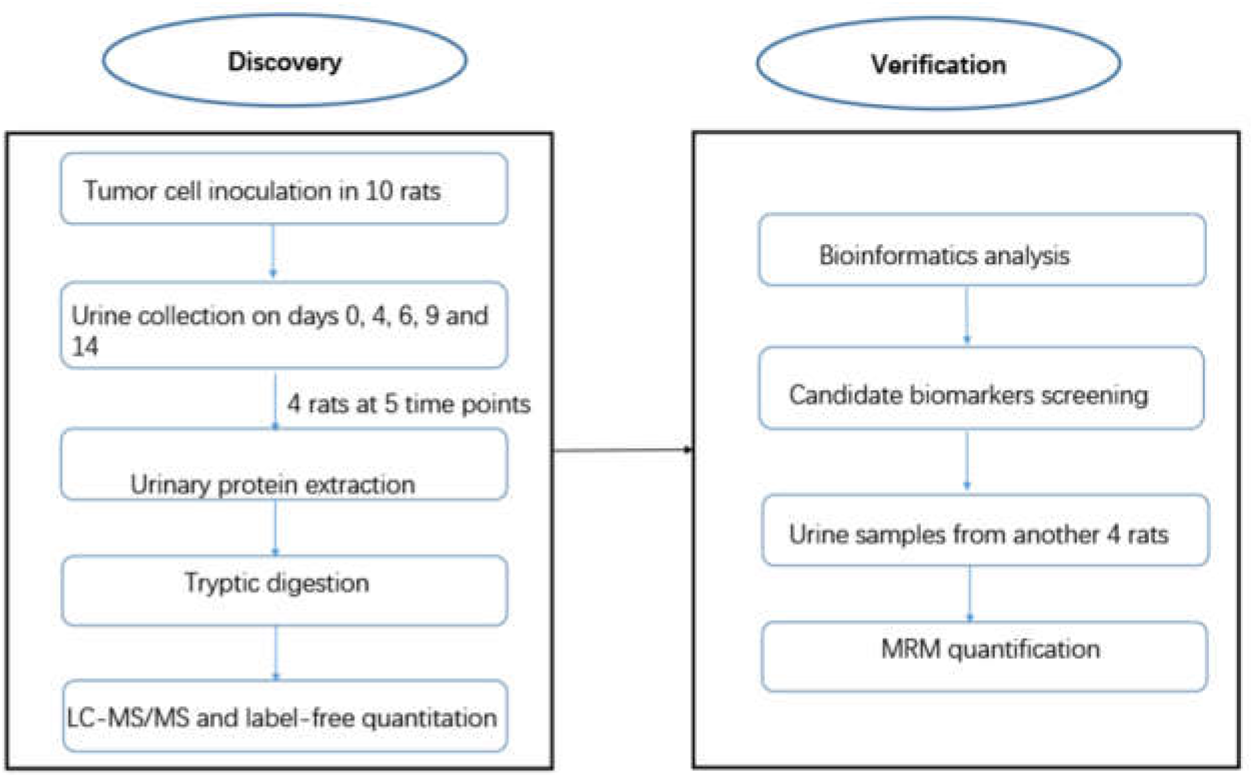
Workflow of urinary proteomics discovery and verification in this study. Urine samples were collected on days 0, 4, 6, 9, and 14 after Walker 256 cell inoculation, and the urinary proteome was analyzed using liquid chromatography coupled with tandem mass spectrometry (LC-MS/MS) identification. Some candidate tumor biomarkers dynamically changed with tumor progression and were verified by multiple reaction monitoring (MRM).

### Unsupervised clustering of the urinary proteome identified at different cancer stages

A biological heat map of clusters from different tumor stages of tumor-bearing rats was produced with the R language (**Fig 4A**). After unsupervised clustering analysis of all urinary proteins identified, it was found that samples at each tumor stage were almost clustered together. The urine samples on day 9 were significantly different from samples at other time points. This result was consistent with the significantly changed protein patterns on SDS-PAGE. This stage represents the phase of rapid tumor growth.

**Fig 4.**
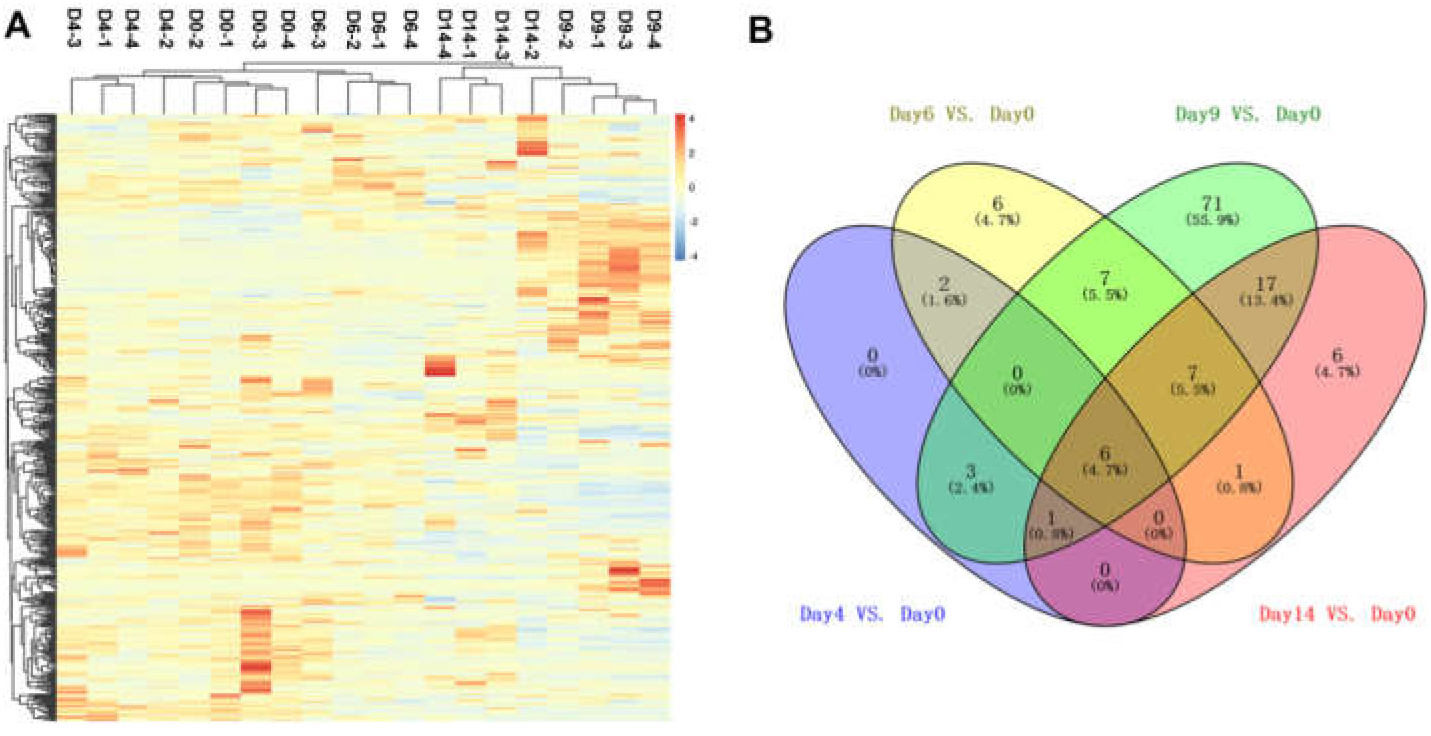
Proteomic analysis of the urine samples of tumor-bearing rats at different phases. (A) Cluster analysis of the proteins identified by LC-MS/MS. (B) Overlap evaluation of the differential proteins identified at different tumor phases.

### Changes in urinary proteomics during tumor progression

The criteria for screening differentially expressed proteins were a fold change ≥ 1.5 compared with day 0 and a p-value < 0.05. Meanwhile, protein spectral counts from all rats in the high-abundance group had to be greater than those in the low-abundance group, and the average spectral count in the high-abundance group had to be more than 3. The details of the differential proteins are shown in **S2 Table**. The overlap of differential proteins identified at different tumor stages is shown by a Venn diagram (**Fig 4B**). There were 12, 29, 112, and 38 differential proteins on days 4, 6, 9, and 14 after W256 cell implantation, corresponding to before the tumor mass was palpable, tumor mass appearance, rapid tumor growth, and cachexia, respectively.

As indicated by the results, many proteins were commonly identified at different time points. Twelve differential proteins (Galectin-3-binding protein, Complement C4, Beta-2-microglobulin, Haptoglobin, Macrophage colony-stimulating factor 1, Coagulation factor XII, and Apolipoprotein A-IV) identified before tumor mass appearance on day 4 were also differentially expressed at later time points. Importantly, six proteins (Haptoglobin, Apolipoprotein A-IV, Complement C4, Beta-2-microglobulin, Alpha-1-acid glycoprotein and Dipeptidyl peptidase 1) were dynamically changed during tumor progression with a fold change >1.5 and p < 0.05, suggesting the potential for these urine proteins to be used for the early detection of tumors. Additionally, urine samples on day 9 had the greatest number of differential proteins (112 proteins), suggesting that the systemic response to tumors in the body is most intense at this stage. This result was consistent with the significantly changed protein patterns from SDS-PAGE analysis. Proteomic changes in urine were probably mediated by factors produced by the tumor and the host response to the presence of the tumor.

### Ingenuity analysis of differential urine proteins in tumor-bearing rats

The biomarker filter function in IPA software was used to filter the candidate cancer biomarkers. Twenty-four differential proteins in this experiment were identified as cancer biomarkers (**S3 Table**). Because W256 cells are breast carcinoma-derived tumor cells, we also screened for breast cancer-related proteins. A total of 27 proteins were associated with breast cancer in previous studies, and 9 of them were identified as biomarkers of breast cancer by the IPA software, namely, ALPL, APOE, CP, CTSC, FABP7, GSTO1, ORM1, PRDX5, and TTR (**S4 Table**).

To identify the major biological pathways involved with the differential urine proteins, IPA was used for canonical pathway enrichment analysis. The proteomics results from this study demonstrate that several pathways, such as acute phase response signaling, LXR/RXR activation, IL-12 signaling and production in macrophages, production of nitric oxide and reactive oxygen species (ROS) in macrophages, clathrin-medicated endocytosis signaling, IL-6 signaling, and apoptosis, were enriched during tumor progression. These pathways changed significantly at different tumor phases during tumor progression.

## MRM verification

At the biomarker validation phase, 30 differential proteins that changed at multiple tumor stages were selected as potentially more reliable cancer biomarkers and were used for MRM verification. The details of these proteins are listed in **Table 1**. Then, urine samples from another four tumor-bearing rats were randomly selected for MRM validation. A total of 26 proteins were successfully quantified, with the exception of LEG9, FA12, CSF1, and GAS6. Finally, 20 differential proteins were changed at multiple time points by MRM-based quantification (**Fig 5**). As a result, 8 differential proteins showed an overall upregulated trend, including A1AG, B2MG, CO4, HPT, LEG5, LG3BP, NGAL, and VCAM1. Twelve proteins showed an overall downregulated trend during tumor progression, including ANTR1, APOA4, ATRN, CALB1, CATC, CO1A1, CRP, CSPG4, PDC6I, PPBT, TCO2, and VTDB. The expression trends of these corresponding proteins were consistent with the results from label-free quantification.

**Table 1.**
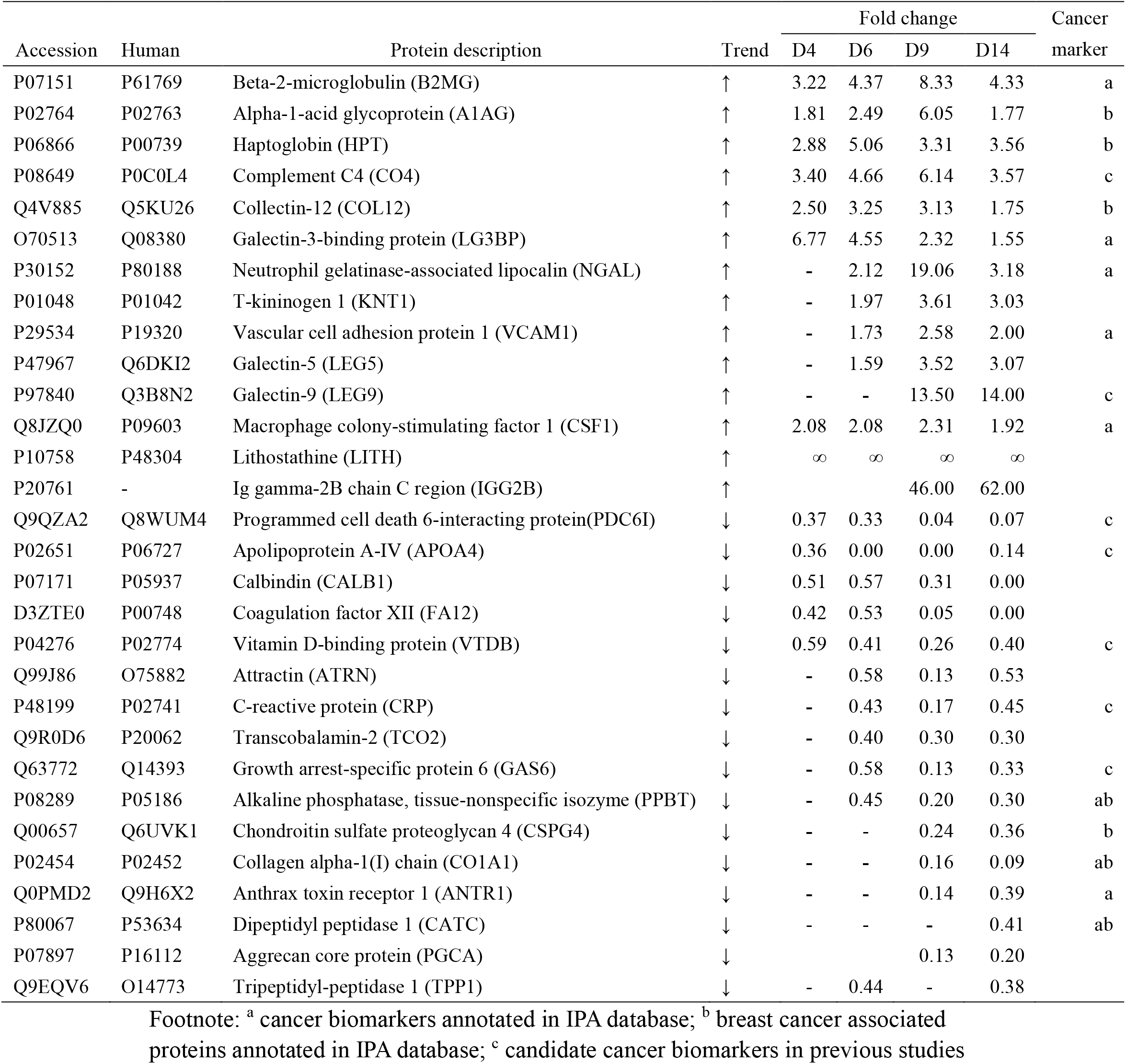
Candidate cancer biomarkers for MRM validation.

**Fig 5.**
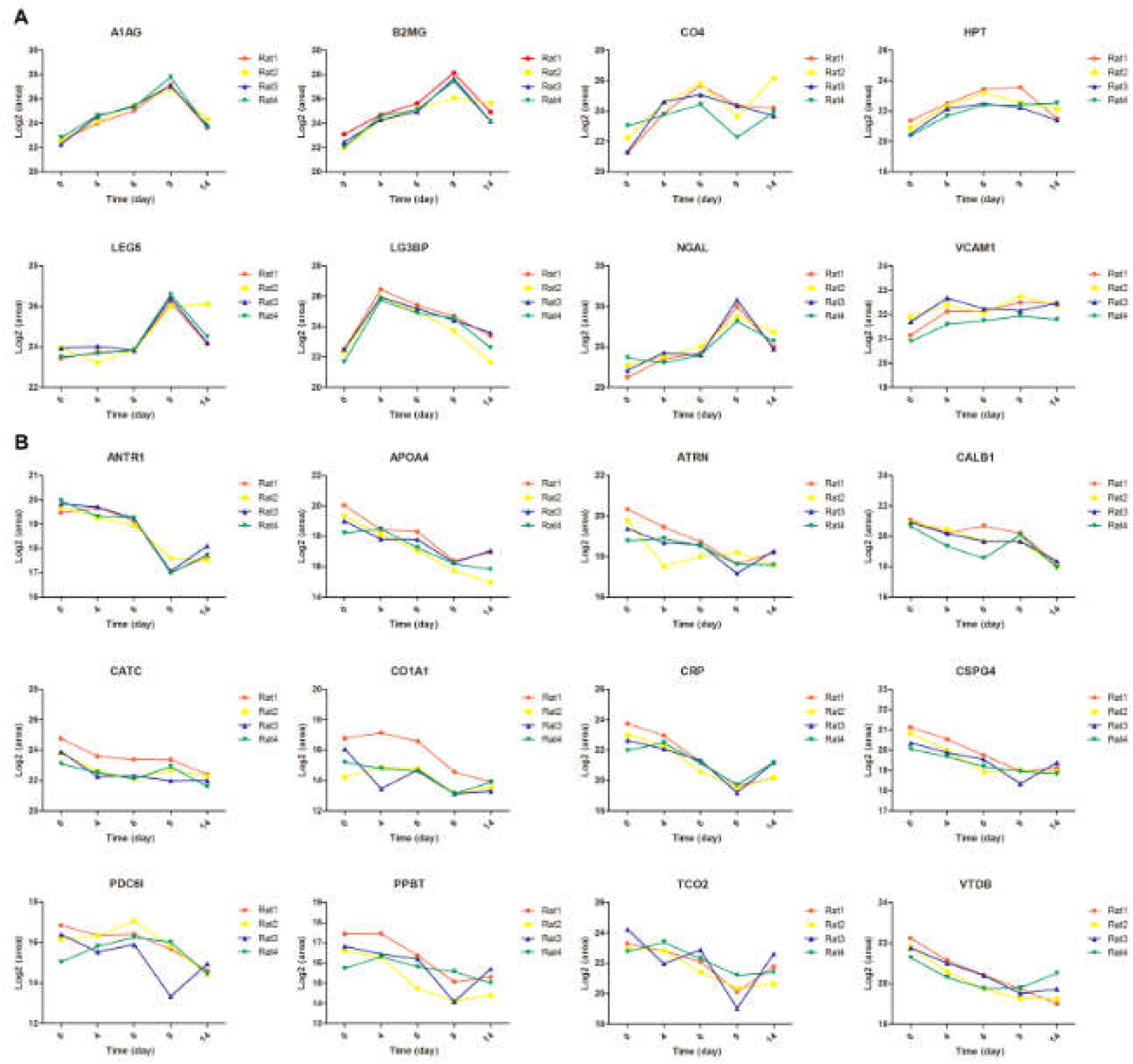
Expression of candidate urine biomarkers from tumor-bearing rats. (A) Eight proteins shared an overall increasing trend in relative abundance. (B) Twelve proteins shared an overall decreasing trend. The x-axis represents different stages after tumor cell inoculation, and the y-axis represents the log_2_ area of intensity based on the protein identified by MRM.

## Discussion

In this study, a tumor-bearing rat model was established by subcutaneous injection of W256 tumor cells. Urine samples were collected from tumor-bearing rats at five time points corresponding to before cancer cell implantation, before the tumor mass was palpable, tumor mass appearance, rapid tumor growth, and cachexia. At the biomarker screening phase, the urinary proteome at different tumor stages was investigated by LC-MS/MS and label-free quantification. The urinary proteome changed significantly with tumor progression. A total of 127 differential proteins were identified at different tumor stages, and 24 proteins were identified as cancer biomarkers, with 9 proteins identified as biomarkers of breast cancer. In a previous study, 408 cancer-associated proteins were reproducibly quantified in urine using targeted proteomics [35]. We transformed differential proteins identified in the urine of tumor-bearing rats to their corresponding human homologous proteins and compared them with the 408 cancer-associated urine proteins. There were 36 common proteins (**S5 Table**). These candidate cancer biomarkers are valuable resources for further clinical verification in the future.

It was interesting that the pathways that changed in this experiment were similar to the pathways that were enriched in a previous urinary proteomics study in breast cancer patients [6]. The pathways included acute phase response signaling, LXR/RXR activation, production of nitric oxide and ROS in macrophages, IL-12 signaling and production in macrophages, and clathrin-medicated endocytosis signaling. Because W256 cells were breast-derived carcinoma cells, it was not a coincidence that there were common changed pathways between the W256 tumor-bearing model and human breast cancer. Meanwhile, 27 differential proteins identified in our experiment were reported as breast cancer-associated proteins.

The Walker-256 carcinoma model has been previously used to study tumor-induced cachexia. It was reported that this model was characterized by reduced food intake and body weight loss starting from day 6 after tumor implantation [36], and the results in our study were consistent with those of previous studies. In our experiment, some metabolic disturbances, such as lipid metabolism and vitamin and mineral metabolism, were observed during tumor progression. The canonical pathway results showed that multiple pathways and networks are involved in the systemic response to tumors. Tumor effects and host responses commonly contribute to these disorders. Proteomics analysis will improve our understanding of the interplay between the tumor and the host at a systemic level. Additionally, cancer cachexia is a multi-organ syndrome, and the main affected tissues include skeletal muscle, liver, heart, fat, and brain tissues [37]. Urine can reveal accumulated systemic changes in the body, thus making urinary proteomics suitable for studying pathophysiological changes during cachexia.

At the biomarker validation phase, to select more reliable cancer biomarkers, 30 differential proteins associated with tumors that changed at multiple time points were further selected for MRM experiments. Ultimately, 26 differential proteins were successfully quantified. The changes in these differential proteins from the MRM results were consistent with corresponding changes in label-free quantification. Further validation in clinical samples is needed.

Several proteins we found were also reported in urine samples of cancer patients. For example, B2MG was found to be a urine marker of several cancers [38]. PDC6I was a potential urinary biomarker of upper gastrointestinal cancer [39]. CO4 in human urine was reported to be helpful in the diagnosis of bladder cancer [40]. NGAL was involved in apoptotic processes. A variety of malignant tumors consistently overexpressed NGAL with increased concentrations in urine, and NGAL was a potential biomarker for malignancy [41]. KNT1 was validated to be a urine marker of breast cancer [42]. Additionally, several proteins were reported to be associated with cancer. For example, LG3BP promotes integrin-mediated cell adhesion and may stimulate host defense against tumor cells. CSF1 promotes reorganization of the actin cytoskeleton and regulates the formation of membrane ruffles, cell adhesion, and cell migration. The association of these 30 urinary proteins with cancer were listed in Table1.

Using label-free quantification, it was found that six proteins (HPT, APOA4, CO4, B2MG, A1AG and CATC) were dynamically changed during tumor progression. Using MRM-based quantification, it was found that in addition to these six proteins, VCAM1, CALB1, CSPG4, and VTDB were also dynamically changed with tumor progression. These 10 proteins were significantly changed even on day 4 after tumor cell implantation and continued their corresponding trends during tumor progression, suggesting the potential of these urine proteins as early cancer markers and for tumor progression monitoring. These urinary proteins were changed even before a tumor mass was palpable. At this stage, the body weights of tumor-bearing rats were not obviously reduced, and the size of the tumor mass might not be detected by imaging equipment. Moreover, the changes in these candidate biomarkers were similar between label-free quantification and MRM quantification. The results suggested that these urine biomarkers will be very valuable in the early detection of tumors (**Table 2**).

**Table 2.**
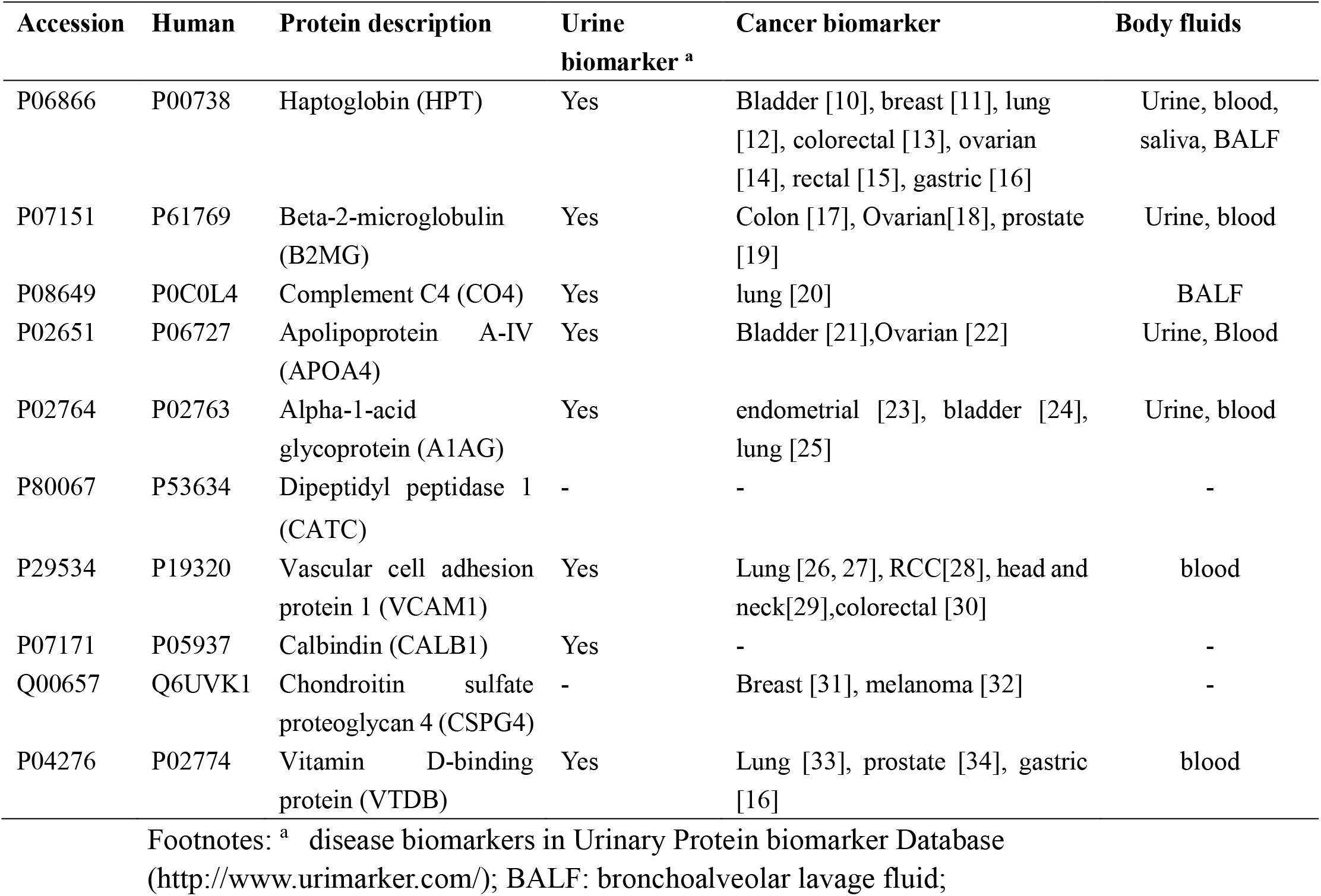
candidate biomarkers for early detection of cancers.

Cancer is a major public health concern, and the early diagnosis of cancer can significantly improve prognosis. There is an urgent need for cancer biomarkers, particularly for early-stage cancer. Although plasma tumor biomarkers are widely used in clinical settings, few have been used effectively for the early detection of cancer, mainly because of their limited sensitivity and/or specificity. Unlike blood, urine lacks mechanisms for maintaining homeostasis; thus, urine is a sensitive biomarker source for detecting pathological conditions, especially the small changes in early stages of diseases. In the current study, urinary proteins provided a preliminary indication of the presence of tumors, even at a very early stage, and some candidate protein biomarkers could be used for early diagnosis and monitoring of cancer progression through urine.

Overall, this study was a preliminary study with a small number of cancer-bearing rats. In future studies, a large number of clinical samples from early-stage cancers are needed to verify the protein patterns of specific biomarkers. The precondition of clinical verification is that urine samples from cancer patients can be collected at an early phase. Urimem, a membrane that can store urinary proteins simply and economically, made large-scale storage of clinical samples possible [43]. The urinary protein biomarkers identified require further evaluation in urine samples of cancer patients to test their sensitivity and specificity for early diagnosis of cancer, and they may also have potential applications in monitoring in cancer treatment and prevention studies.

